# The genetic signatures of *Salmonella* Typhi carriage in the human gallbladder

**DOI:** 10.1101/2020.06.08.140053

**Authors:** Pham Thanh Duy, Nga Tran Vu Thieu, Nguyen Thi Nguyen To, Ho Ngoc Dan Thanh, Sabina Dongol, Abhilasha Karkey, Megan Carey, Buddha Basnyat, Gordon Dougan, Maia A Rabaa, Stephen Baker

## Abstract

Despite recent gains in typhoid fever control, asymptomatic carriage of *Salmonella* Typhi in the gallbladder remains an enigma. Aiming to understand if *S.* Typhi in the gallbladder are vital for transmission and/or adapted for long-term colonisation we performed whole genome sequencing on a collection of *S*. Typhi isolated from the gallbladders of typhoid carriers. These sequences were compared to contemporaneous sequences from organisms isolated from the blood of acute patients.*S*. Typhi carriage was not restricted to any particular genotype or conformation of antimicrobial resistance genes but reflective of the general population. However, gallbladder isolates had a higher genetic variability than acute isolates, with median pairwise SNP distances of 21 and 13 SNPs (*p*=2.8×10^−9^), respectively. This variation was associated with a higher prevalence of nonsense mutations in the gallbladder isolates in the predominant genotype. Notably, gallbladder isolates displayed a higher frequency of non-synonymous mutations in genes encoding hypothetical proteins, membrane lipoproteins, transport/binding proteins, surface antigens, and carbohydrate degradation. Particularly, we identified several gallbladder-specific non-synonymous mutations involved in LPS synthesis and modification, with some isolates lacking the Vi capsular polysaccharide vaccine target due to a 134Kb deletion. Long-term gallbladder carriage of *S*. Typhi results in atypically long branch lengths that can distinguish between carriage and acute infection. Our data strongly suggests typhoid carriers are unlikely to play a principal role in disease transmission in endemic settings, but that the hostile environment of the human gallbladder may generate new antigenic variants through immune selection.

## Background

Typhoid fever, a potentially life-threatening systemic infection caused predominantly by the bacterium *Salmonella enterica* serovar Typhi (*S.* Typhi), remains a significant public health problem in resource-poor settings including parts of Asia and Africa ^1^. The disease is contracted via ingestion of contaminated food or water or through contact with individuals excreting the organism ^2^. The majority of typhoid patients fully recover with appropriate treatment; however, some individuals can become asymptomatic carriers and shed infectious bacteria in their faeces for an ill-defined period of time. Asymptomatic carriage of *S*. Typhi has been recognized as a public health threat for more than a century, with infamous typhoid carriers like Mary Mallon, a cook in New York, and Mr N, a milker in England, identified in the early part of the 20^th^ century ^3,4^.

Typhoid carriage can be differentiated into three categories depending on the duration of shedding: convalescent (three weeks to three months), temporary (three to twelve months), and chronic (more than one year) ^5^. In endemic regions, an estimated 2-5 percent of acute typhoid patients become chronic carriers, meaning that they continue to intermittently shed the bacteria indefinitely after apparent clinical resolution ^3,5^. Consequently, chronic carriers are widely believed to be an ecological niche that facilitates the transmission and persistence of typhoid in human populations ^6^. *S*. Typhi is a human-restricted pathogen, meaning that the disease may be theoretically eliminated locally by reducing transmission through targeted treatment, improved sanitation, and mass vaccination. Consequently, understanding the role of chronic carriers in disease transmission, and detecting them prospectively, may accelerate disease elimination.

Despite substantive gains in understanding the biology of typhoid, we have generated limited new insights into typhoid carriage in recent decades. Data from murine models of *Salmonella* carriage and human clinical investigations have determined that the gallbladder is a key permissive niche for long-term bacterial persistence ^7–13^. Various epidemiological investigations have shown that gallstones and gallbladder damage may facilitate typhoid carriage ^9,13–17^, and that *Salmonella* preferentially attach to, and form biofilms on, cholesterol-rich gallstones ^7,11,13,18,19^. *S*. Typhi carriage isolates have been previously genetically compared with isolates from acute infection, with the aim of identifying signatures associated with carriage ^20–23^. However, these studies were unable to infer how carriage isolates directly relate to those causing acute disease.

It is apparent we need a better understanding of the role of the typhoid carrier and associated organisms to generate new approaches to the management of such individuals in endemic locations. Although it is widely accepted that that *S*. Typhi carriage play a key role in the transmission of typhoid in endemic settings it is unknown if carriage organisms are somehow adapted for long-term colonisation. Aiming to address this question, we performed whole genome sequencing and detailed genetic analyses on *S*. Typhi isolated from the gallbladders of typhoid carriers in Kathmandu. We compared these data to the sequences of contemporaneous organisms isolated from the blood of acutely infected patients in the same community over the same time period. Our data provides new insight into the role of typhoid carriage in disease transmission, showing that whilst carriage isolates are reflective of the general *S*. Typhi population circulating in the community, gallbladder carriage subjects organisms to immune pressures, which induces genetic variation and genomic degradation.

## Results

### The phylogenetic relationships between acute and gallbladder S. Typhi isolates

Between June 2007 and October 2010, we conducted a *Salmonella* carriage study in Kathmandu^13^. Patients undergoing cholecystectomy for acute or chronic cholecystitis were enrolled; bile and stool samples from 1,377 individuals were collected and subjected to microbiological examination. Twenty-four *S*. Typhi were isolated from bile samples taken from these patients and designated as gallbladder isolates. Ninety-six *S*. Typhi isolates recovered from patients with acute typhoid fever living in the same population over the same time period were used for comparison ^24^ (denoted as acute isolates) (Table S1). A phylogenetic analysis of these 120 *S*. Typhi isolates demonstrated that subclade 4.3.1 (H58) was the dominant genotype, constituting 62.5% (15/24) of all gallbladder isolates and 65.6% (63/96) of all acute isolates. The second most common genotype was 3.3.0 (H1), accounting for 12.5% (3/24) and 14.6% (14/96) of all gallbladder and acute isolates, respectively.

We identified a significant degree of genetic diversity within this collection of acute and carriage organisms, with multiple less-common genotypes co-circulating, included various clades (4.1, 3.1 and 2.2), subclades (3.2.2, 3.0.1, 2.2.2 and 2.2.1), and organisms within primary cluster 2 (Figure 1). The less common genotypes from the gallbladder fell within subclade 3.2.2 (8.3%; 2/24), 2.2.2 (4.2%; 1/24), clade 2.2 (8.3%; 2/24) and primary cluster 2 (4.2%; 1/24). Overall, gallbladder isolates were not significantly associated with subclade 4.3.1 in comparison with other genotypes (15/24 versus 9/24, *p*=0.083; Chi-squared test). These initial observations indicate that *S*. Typhi carriage was not restricted to any particular *S*. Typhi genotype; instead, the genotype distribution among gallbladder isolates generally reflected a genetic structure similar to that of the acute *S*. Typhi infections circulating in the community.

**Figure 1.**
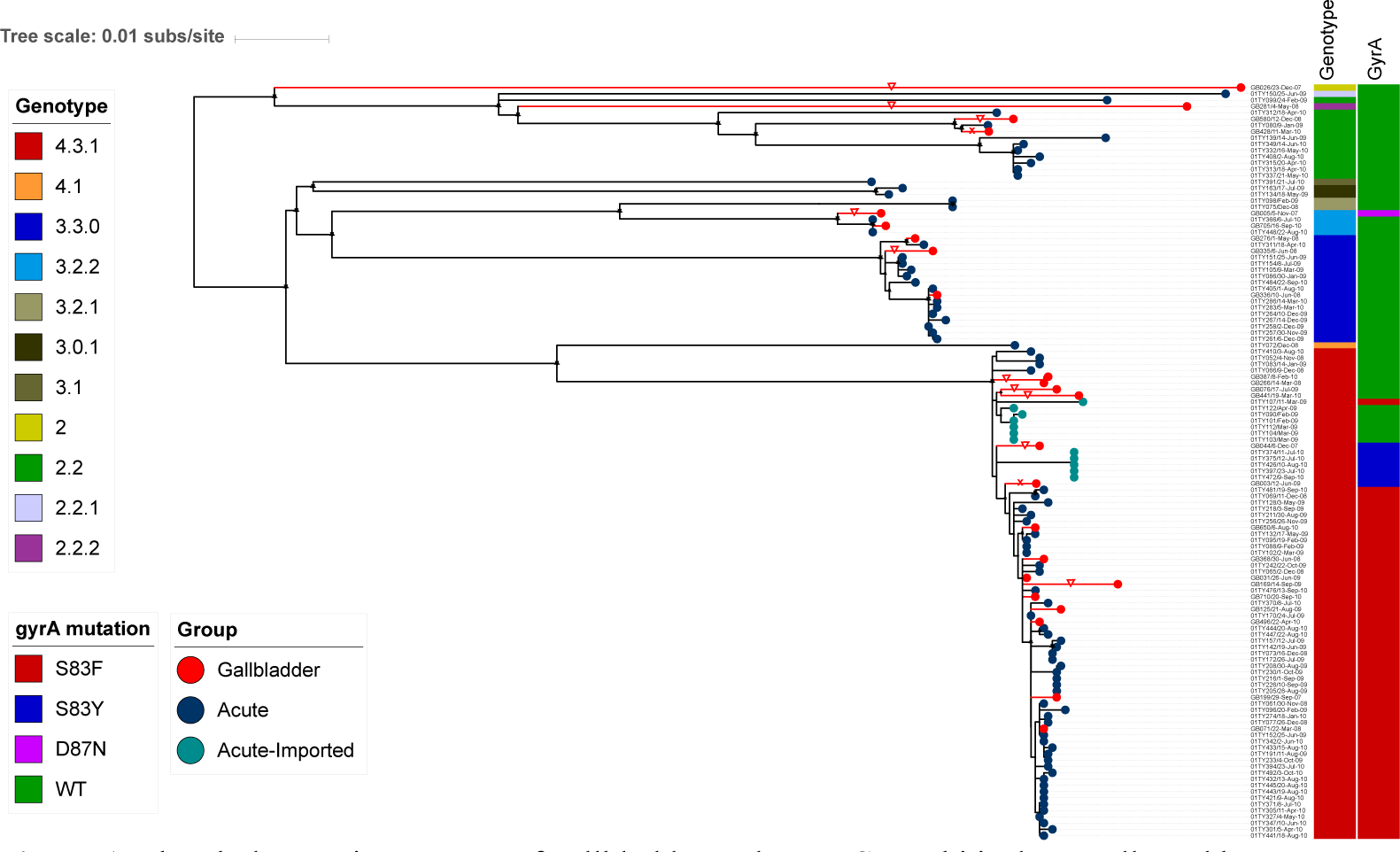
The phylogenetic structure of gallbladder and acute *S*. Typhi isolates collected between 2007 and 2010. Rooted maximum likelihood tree (*S*. Paratyphi A used as an outgroup to root the tree and pruned for visualization) based on core-genome SNPs of 120 *S.* Typhi isolates with the corresponding metadata: genotype, *gyrA* mutation. Gallbladder and acute isolates are shown as red and dark circles at the terminal nodes, respectively. Acute isolates originating from importation are also highlighted by turquoise circles at the terminal nodes. Terminal branches leading to gallbladder isolates are highlighted in red. Red triangles show gallbladder isolates associated with unusually long terminal branches.

### Antimicrobial susceptibility

We speculated resistance to key antimicrobials may facilitate the development of carriage. However, we found that the *S*. Typhi gallbladder isolates did not carry any obviously acquired AMR genes. However, chromosomal mutations associated with reduced susceptibility against fluoroquinolones were common. These fluoroquinolone resistance-associated mutations within the gallbladder organisms were more commonly observed in subclade 4.3.1 than in non-subclade 4.3.1 (73% (11/15) versus 11% (1/9), *p*=0.01; Chi-squared test). In comparing the respective *gyrA* mutation profiles between the acute and gallbladder isolates within subclade 4.3.1, we found that 76.2% (48/63) and 60% (9/15) carried the S83F mutation respectively, 7.9% (5/63) and 13.3% (2/15) carried the S83Y mutation, and 15.9% (10/63) and 26.7% (4/15) had no *gyrA* mutation. Consequently, there was no significant difference (*p=*0.327; Chi-squared test) in the presence of fluoroquinolone resistance-associated mutations between acute and gallbladder isolates within subclade 4.3.1.

### Phylogenetic signatures of long-term Salmonella Typhi carriage

Despite the acute and gallbladder *S*. Typhi isolates generally clustering within the same genotypes across the phylogeny, we observed that a substantial proportion of the gallbladder isolates had higher genetic variability, which could be distinguished by long terminal branches (Figure 1). The median pairwise SNP distance of gallbladder isolates within subclade 4.3.1 was 21 SNPs (IQR: 12-24), which was significantly greater than that of the corresponding acute isolates (13 SNPs (IQR: 8-19 SNPs) (*p*=2.8×10^−9^, Wilcoxon rank-sum test) (Figure S2). Similarly, the median pairwise SNP distance of gallbladder isolates within subclade 3.3.0 (20 SNPs, IQR: 13-22 SNPs) was higher than that of acute isolates (13 SNPs, IQR: 4-15 SNPs) (*p*=0.26, Wilcoxon rank-sum test).

We mapped the contemporary acute and gallbladder *S*. Typhi sequences onto the global *S*. Typhi phylogeny, which indicated that the majority of these Nepalese acute and gallbladder *S*. Typhi isolates fell within known domestic genotypes, with limited evidence of importation from other countries (Figure S1). This observation suggests that the long terminal branches associated with gallbladder isolates were unlikely to be driven by the importation of these organisms from alternative countries.

We next estimated and plotted the nearest phylogenetic distances (NPDs) between each taxon and its nearest neighbour on the subclade 4.3.1 tree versus the year of isolation, the age of the individual from whom the organism was isolated, and the *gyrA* mutation profile. We hypothesized that the annual distribution of NPDs of *S*. Typhi acute isolates would represent the phylogenetic diversity (mutation accumulation) occurring annually via acute disease transmission and would be comparable over multiple years. Alternatively, we considered that *S*. Typhi in the gallbladder may develop characteristic adaptive mutations facilitating long-term persistence, causing them to gradually become increasingly distinct from contemporaneous acute isolates, leading to greater phylogenetic distances relative to their neighbours. In addition, given that all acute subclade 4.3.1 isolates here exhibited a *gyrA* mutation, the gallbladder subclade 4.3.1 isolates without a *gyrA* mutation were more likely to have colonized the gallbladder prior to nalidixic acid resistance becoming commonplace.

**Figure S1.**
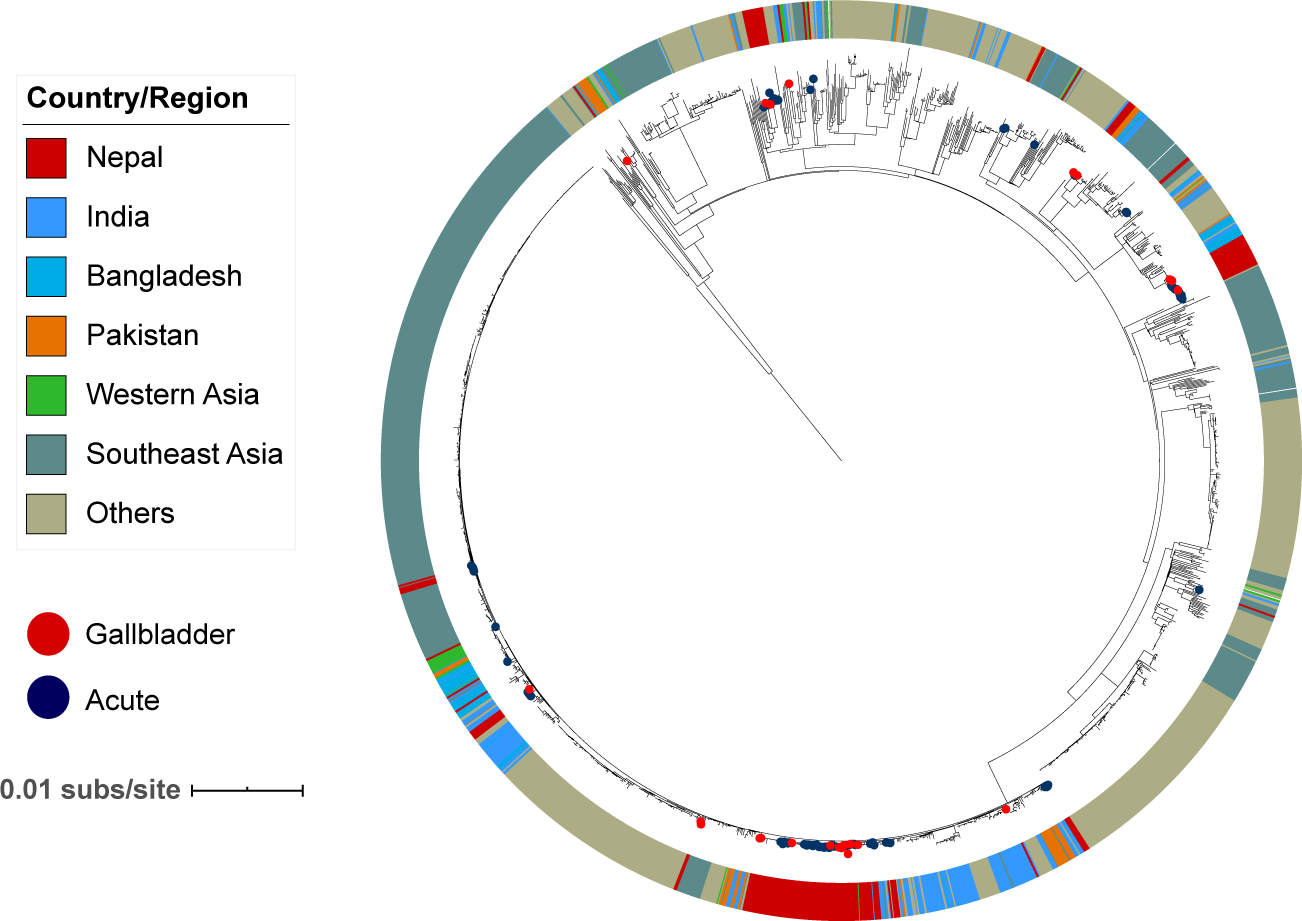
Phylogenetic structure of acute and gallbladder *Salmonella* Typhi isolates from Nepal in the global context. Acute and gallbladder *S.* Typhi isolates from this study are highlighted in blue and red circles; respectively, at the terminal nodes. The outer ring exhibits the location of the isolates from Nepal and its neighbouring countries as well as other regions in the world.

Our analyses showed that the average (±SD) NPD per year of acute subclade 4.3.1 isolates was comparable; specifically, 0.00163 (± 0.00202) substitutions/site (∼3.6 (± 4.4) SNPs) in 2008; 0.00110 (± 0.00229) substitutions/site (∼2.4 (± 5) SNPs) in 2009, and 0.00144 (± 0.00238) substitutions/site (∼3.2 (± 5.2) SNPs) in 2010. The majority of the subclade 4.3.1 gallbladder isolates (8/10) for which NPDs fell within the annual NPD distribution of acute subclade 4.3.1 isolates were associated with comparable terminal branch lengths and had a *gyrA* S83F mutation. Based on these findings, we surmised that gallbladder colonization with these isolates was likely to have occurred relatively recently in these individuals. Notable exceptions were two gallbladder isolates (GB266 and GB387) that did not possess a *gyrA* mutation and were associated with long terminal branches but had low NPDs as they clustered closely within the main phylogeny (Figures 1 and 2). Further, our data showed that all subclade 4.3.1 gallbladder isolates exhibiting abnormally high NPDs were associated with long terminal branches, indicative of chronic carriage (Figure 2). In particular, two subclade 4.3.1 gallbladder isolates (GB76 and GB441) lacked *gyrA* mutations, two others (GB003 and GB044) had *gyrA* S83Y mutations, and the remaining one (GB169) exhibited *gyrA* mutation S83F. With respect to the age distribution, typhoid carriers were significantly older (median age 36 years, range: 20-67) than patients with acute illness (median age 16 years, range: 0-31) (*p*=3.8×10^−8^, Wilcoxon rank-sum test). The gallbladder isolates thought to have originated from chronic carriers (based on above data) were obtained from individuals between aged between 27 and 40 years, which was older than the majority of the sampled acute typhoid patients; however, there was no significant difference in age distribution between those estimated to be recent and chronic carriers (Figure 2).

**Figure 2.**
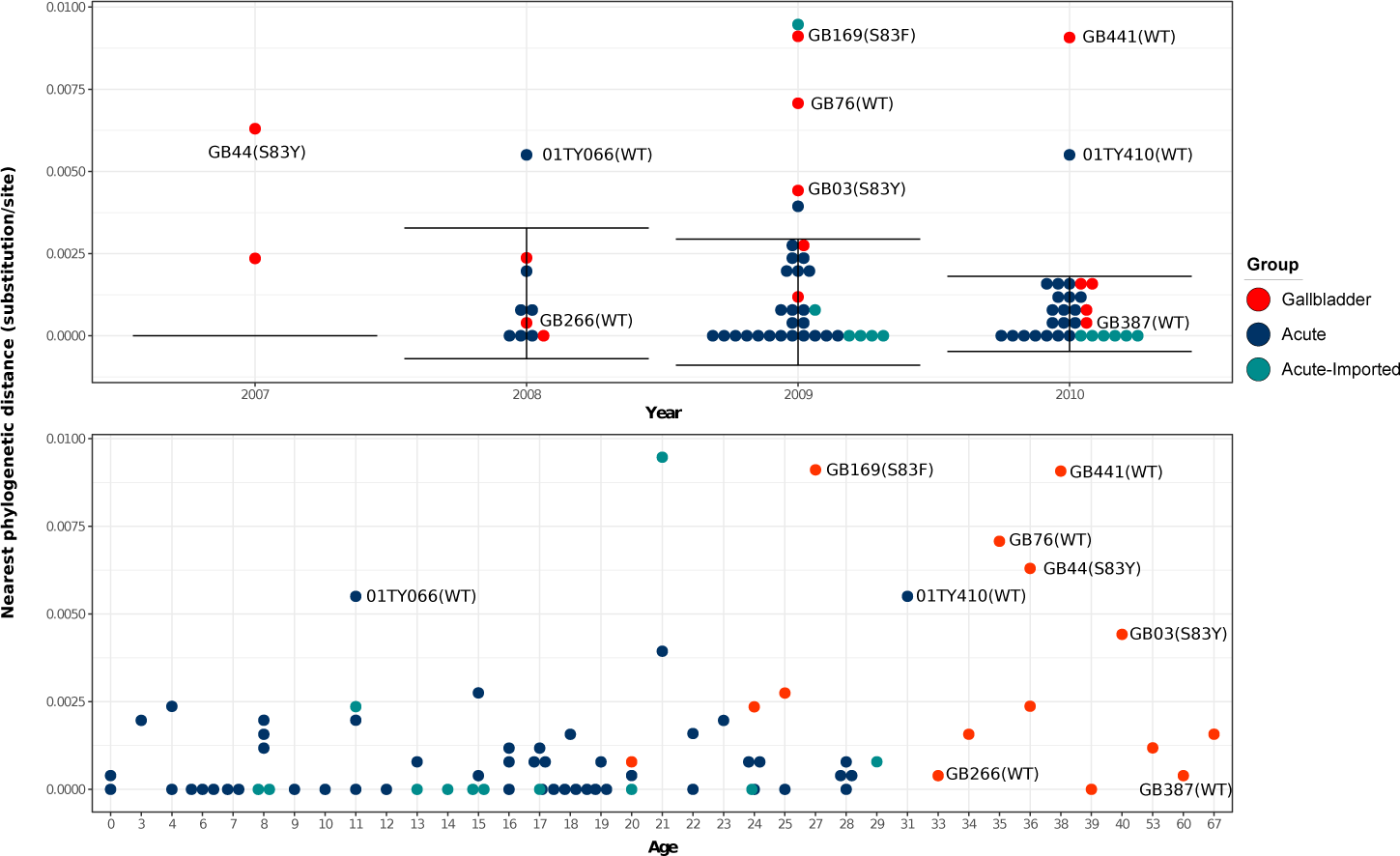
The Distribution of nearest phylogenetic distances of gallbladder and acute H58 isolates over the study period. Each circle represents the phylogenetic distance from each isolate to its nearest neighbour on the phylogenetic tree. The error bar represents the average phylogenetic distance to the nearest neighbour (± standard deviation) for acute H58 isolates. Gallbladder and acute isolates estimated to have originated from chronic carriers are labelled with their corresponding strain names.

### The genetic traits of Salmonella Typhi gallbladder isolates

To identify potentially adaptive mutations associated with typhoid carriage, all nonsynonymous SNPs (NSs) occurring exclusively within the *S*. Typhi gallbladder genome sequences were identified and grouped by their predicted or known function. A corresponding analysis was performed for all NSs in the acute *S*. Typhi isolates. A total of 228 gallbladder-specific NSs (212 missense and 16 nonsense mutations) and 469 acute-specific NSs (437 missense and 32 nonsense mutations) were identified. In general, there was no significant difference (*p*=0.924; Chi-square test) in the proportion of nonsense mutations out of total specific NSs in the gallbladder versus the acute isolates across all genotypes. However, among subclade 4.3.1 isolates, the proportion of nonsense mutations out of total specific NSs was significantly higher for gallbladder isolates than for acute isolates (10/60 compared to 2/67, Fisher exact test, *p*=0.01). These data suggest that gene degradation resulting from nonsense mutations was more common in the subclade 4.3.1 gallbladder isolates compared to the subclade 4.3.1 acute isolates.

Inactivated genes in the gallbladder isolates included genes involved in the synthesis of peptidoglycan (*pbpC*), vitamin B12 receptor (*btuB*), general stress response regulator (*rpoS*), a laterally acquired protein in SPI-7 (STY4562), membrane transport protein (STY3932), central metabolism (STY0230, *ggt*), hypothetical proteins (STY0929 and STY4178), and osmotically inducible lipoprotein E precursor (*osmE*) (Table 1).

**Table 1.**
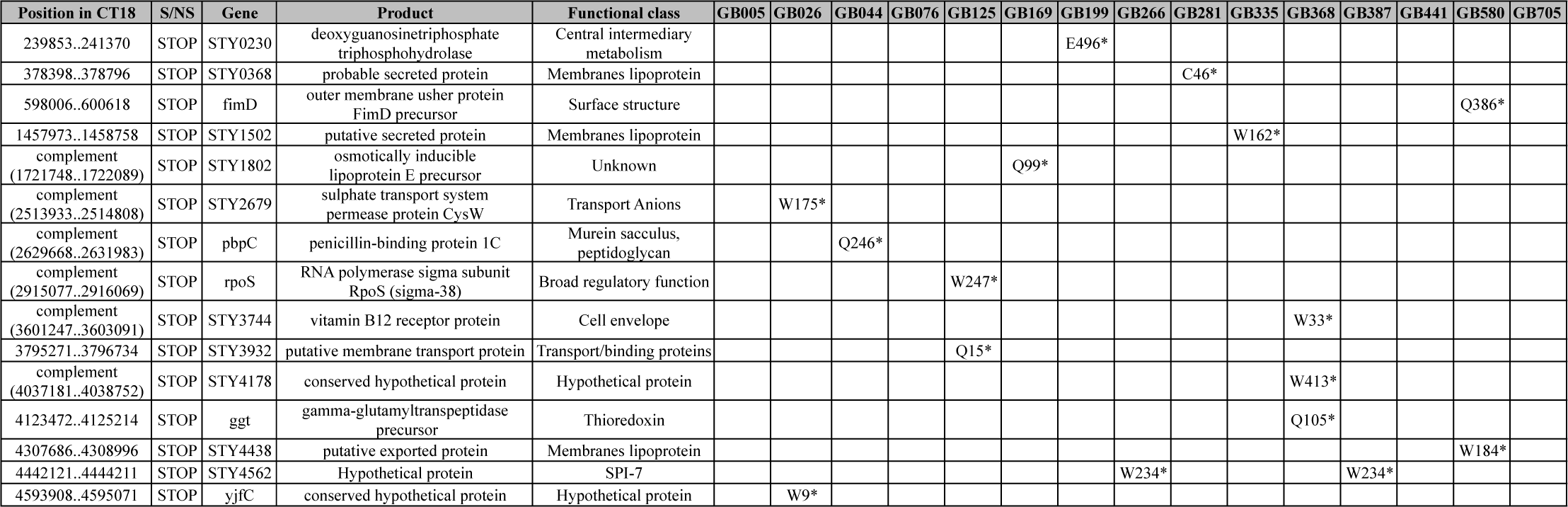
Nonsense mutations and their predicted functions in gallbladder isolates

Overall, the gallbladder- and acute-specific NSs across all genotypes could be grouped into 78 functional categories. The highest prevalence of these NSs was found in genes encoding hypothetical proteins, membrane lipoproteins, unknown functions, transport/binding proteins, SPI-7, general regulatory functions, surface polysaccharides and antigens, carbohydrate degradation, and DNA replication/modification. The proportions of NSs in SPI-7, surface polysaccharides and antigens, pathogenicity, cell envelope, anaerobic respiration, fatty acid biosynthesis and transport/binding proteins were higher in gallbladder than acute isolates (Figure 3). Notably, the data showed that the proportion of NSs in the *viaB* operon (encoding the Vi antigen, target of the typhoid conjugate vaccine (TCV)) was significantly higher in gallbladder isolates compared to the acute isolates across all genotypes (9/228 compared to 7/469, Chi-squared test, *p*=0.04). Similar results were obtained when considering only *S*. Typhi isolates belonging to subclade 4.3.1, with gallbladder isolates having more specific NSs in the *viaB* operon than the acute isolates (5/60 compared to 1/67, Fisher’s exact test, *p*=0.09). Additionally, we identified two gallbladder isolates (GB428 and GB003) that had lost the Vi capsular polysaccharide due to the deletion of the entire SPI-7 region (134kb).

**Figure 3.**
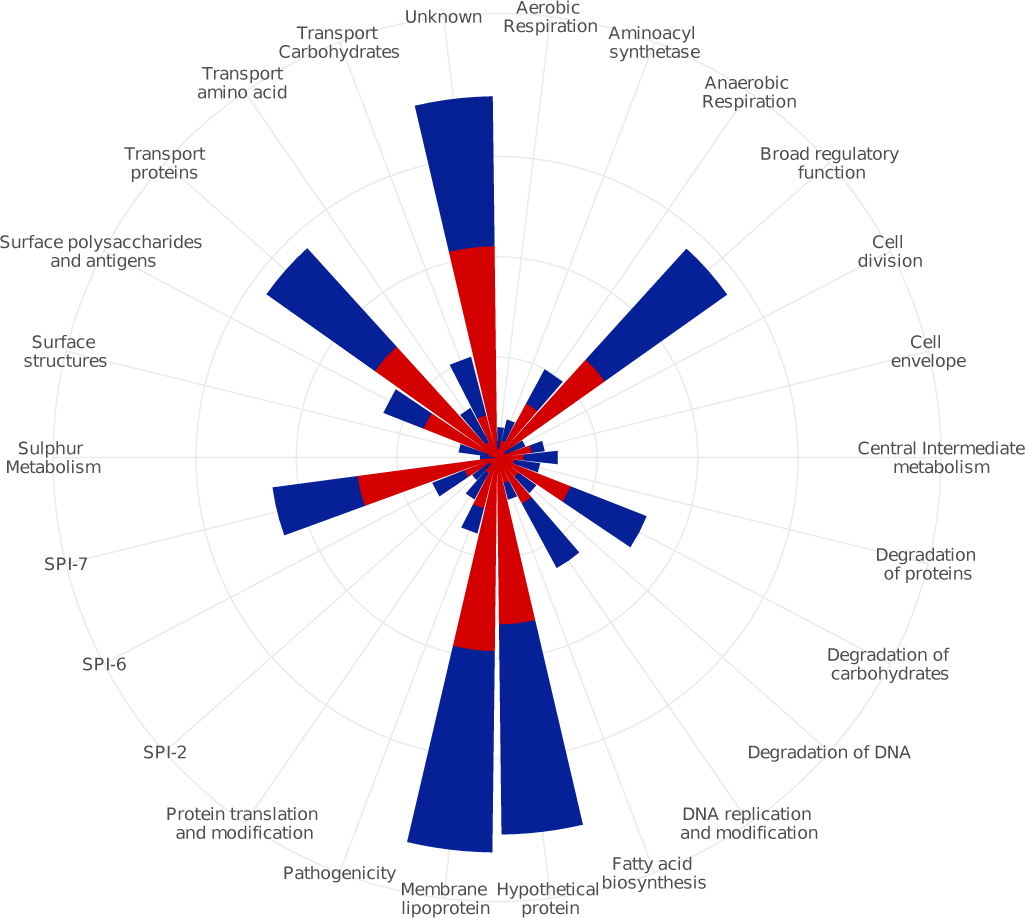
Functional classes of *Salmonella* Typhi genes associated with the highest prevalence of gallbladder-specific nonsynonymous SNPs versus acute-specific nonsynonymous SNPs Functional classes are shown on the outermost circle. Four circles from the middle represent 5 – 10 – 15 – 20 percent of the cumulative percentage of functional classes. Red and blue blocks are representatives of gallbladder and acute isolates, respectively.

### Positive selection associated with typhoid carriage

Finally, we investigated signatures of positive selection by identifying analogous genetic variation detected in different gallbladder isolates. Among the gallbladder specific NSs, a number of different mutations were present in the same gene or the same biological pathway in at least two phylogenetically unlinked gallbladder isolates. For example, within the *viaB* operon, there were two NSs at codon 137 and 462 in the *tviE* gene (isolates GB580 and GB026) and six NSs in codons 166, 504, 506, 508, 665 and 752 in the *tviD* gene (isolates GB005, GB026, GB076, GB125 and GB281, respectively). Both genes facilitate the polymerization and translocation of the Vi capsule ^25^. Convergent NSs were also observed in the *rpoS* gene (sigma factor sigma-38) of isolates GB125 (nonsense mutation at codon 247) and GB705 (NSs at codon 94 and 250), which may impact general stress response and nutrient starvation. A further example was NSs at codon 59 and 230 in the *degS* gene (serine protease) (isolates GB005 and GB169). *DegS* is a component of the DegS-DegU two-component regulation system involved in expression of several degradative enzymes for salt stress responses and growth-limiting conditions. Additionally, three isolates (GB005, GB026 and GB705) each possessed an NS (codons 335, 406 and 946, respectively) in STY1242 (*ptsG* - glucose-specific PTS system IIBC component). PtsG enzyme is a component of the glucose-specific phosphotransferase system and plays a role in phosphorylation and translocation of glucose across the bacterial membrane, and is induced in carbon-limited conditions ^26^. NSs in several other genes were observed in >2 carriage isolates, including STY0429 (*SbcC* - exonuclease), STY0661 (*dmsC* - molybdopterin containing oxidoreductase membrane anchor subunit), STY1447 (putative ribulose-5-phosphate 3-epimerase) and STY2760 (*ratA* - putative exported protein) (Table 2).

**Table 2.**
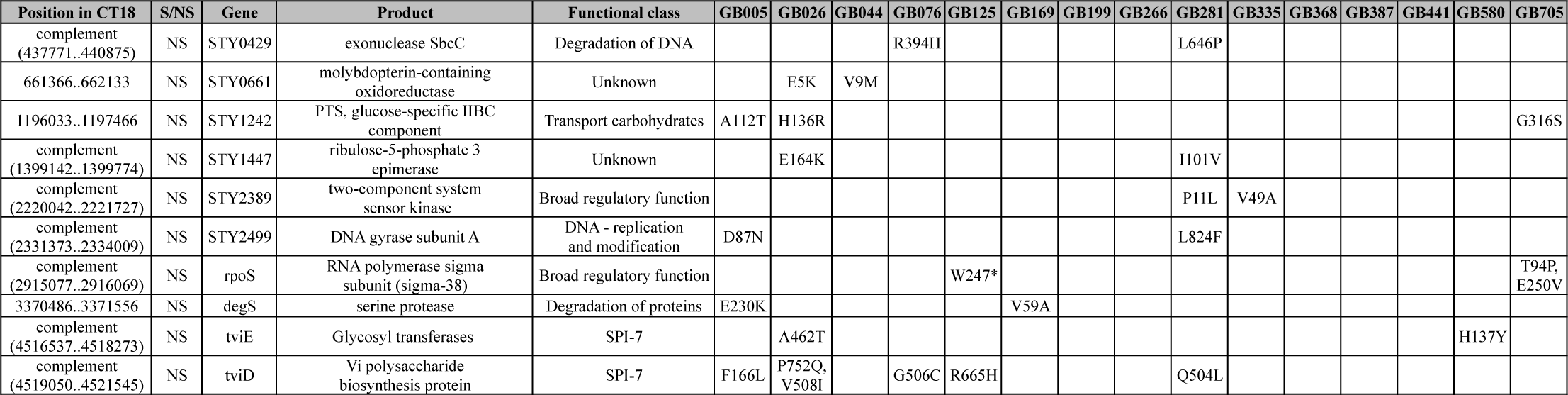
Nonsynonymous mutations associated with positive selection in gallbladder isolates

### Evidence of selective pressure on lipopolysaccharide

With respect to convergent mutations within the same biological pathways, there were a number of gallbladder-specific NSs involved in LPS O-antigen synthesis and modification; for example, an NS in the *rfc* gene (regulator of O-antigen polymerization) in isolate GB441, an NS in STY2629 (LPS modification acyltransferase) in isolate GB335, two NSs in *rfbE* (CDP-tyvelose-2-epimerase) and *rfaG* genes (LPS core biosynthesis protein) in isolate GB281, and three NSs in the *rfbK* (phosphomannomutase), *manB* (phosphomannomutase), and *rfaD* genes (ADP-L-Glycero-D-mannoheptose-6-epimease) in isolate GB026. *RfbK* and *manB* are both related to GDP-mannose synthesis for the LPS, and *rfaD* is an enzyme that catalyzes the conversion of ADP-D-glycerol-D-mannoheptose to ADP-L-glycerol-D-mannoheptose, a precursor for the synthesis of inner-core LPS.

## Discussion

As stakeholders consider introduction of a new TCV into their national immunization programmes, research into the role of chronic carriers in bacterial persistence and disease transmission in endemic settings is needed to forecast the longer-term impact of vaccination on the transmission dynamics of typhoid and inform appropriate public health measures. However, epidemiological investigations of typhoid carriage are challenging, given that this population is problematic to identify and follow. Currently, the environmental factors driving the evolution of *S.* Typhi within the gallbladder are poorly understood and little is known about the adaptive mechanisms that may promote long-term survival. Our study is the largest genomic investigation of *S*. Typhi gallbladder carriage in a typhoid endemic setting, which allowed us to provide unprecedented insight into the genetic and phylogenetic signatures associated with typhoid carriage, but also to utilize these data to infer the potential role of typhoid carriage in disease transmission.

**S1 Table.** Gallbladder and acute *Salmonella* Typhi isolates and their associated metadata

Our data demonstrated, contrary to previous suggestions ^27^, that carriage of typhoid in the gallbladder was not restricted to any particular genotype and was associated a diverse range of bacterial genotypes, which largely mirrored the genetic structure of the bacterial population causing acute disease in Nepal. Further, typhoid carriage was not confined to specific AMR phenotypes, signifying that carriage is not associated with treatment failure with specific antimicrobials interacting with corresponding AMR profiles. However, by comparing the pairwise SNP distances between gallbladder and acute isolates within the same genotype, we found that gallbladder isolates displayed significantly greater genetic diversity compared to acute isolates, which suggests that long-term exposure to the gallbladder environment results in different accumulated adaptive mutations over time than would be generated in acute isolates. Our phylogenetic reconstruction of *S.* Typhi revealed that a number of gallbladder isolates had atypically long terminal branches, signifying that chronic carriage isolates may have a distinct phylogenetic signature which could be potentially utilized for the identification of organisms arising from chronic carriers. Further investigating this phenomenon, we found that the annual distribution of NPDs of acute isolates, which likely reflects mutation accumulation in the natural environment, was highly comparable across years and could be exploited to disaggregate recent carriers from longer-term carriers. If carriers are relevant, then we would predict they would be proportionally more important in causing acute disease in immunised populations with reduced environmental transmission. Therefore, we can use the annual NPD distribution to assess the impact of typhoid vaccination on disease transmission dynamics in endemic areas.

The role of chronic carriage in disease transmission represents one of the most long-standing questions in typhoid fever. Though typhoid carriers have been widely considered as an important source of infection, their exact contribution to transmission in endemic areas is not well understood. Previous molecular epidemiological studies in endemic regions highlighted an abundance of long-cycle environmental transmission in these settings, with a wide diversity of co-circulating bacterial genotypes isolated from acute typhoid patients ^28–31^, suggesting that person-to-person transmission makes a minimal contribution to new typhoid cases in an endemic area. Here, few gallbladder isolates clustered in close proximity or were directly linked with acute isolates and had long terminal branches. These observations suggest that these organisms play a negligible role in causing onward acute infections. Notably, none of the pre-surgical stool cultures from these patients undergoing cholecystectomy were positive for *S*. Typhi. However, the infectivity and transmission fitness of gallbladder isolates must be investigated further, as we cannot rule out the possibility that gallbladder isolates can become a more important source of infection when environmental transmission is successfully reduced. Further, the fact that gallbladder isolates display greater genetic variation than acute isolates implies that the gallbladder may act as an important ecological niche for generating novel genotypes.

By identifying NS mutations occurring specifically in gallbladder isolates and classifying them into predicted functional classes for comparison with those of acute isolates, we found that gene degradation by nonsense mutation was significantly higher in gallbladder compared to acute isolates within subclade 4.3.1. The effects of gene inactivation on phenotype, fitness and adaptation of carriage isolates inside the gallbladder are currently unknown. Further investigation of this phenomenon is necessary, as gene inactivation has been shown to be an important molecular mechanism in human adaptation in the evolutionary history of *S.* Typhi ^32,33^.

We additionally found evidence for the enrichment of NSs in genes encoding the Vi polysaccharide capsule in gallbladder isolates. The Vi antigen is immunogenic and anti-Vi antibody gradually wanes in acute typhoid patients after recovery, but can be detected in plasma from chronic carriers ^34,35^. Data from sero-surveillance studies for chronic carriage have commonly reported elevated anti-Vi antibodies in healthy individuals, which could be associated with carriage or repeated infections ^36,37^. Immunofluorescent staining of biofilms produced by *S*. Typhi on the surface of human gallstones demonstrated an abundance of Vi capsule on the surface of the colonising bacteria, suggesting that *S*. Typhi constitutively expresses Vi during carriage ^19^. The increased frequency of nonsynonymous mutations in the viaB operon (*tviB, tviD* and *tviE*) of gallbladder isolates, combined with high anti-Vi antibody titres in plasma ^38^ suggest that *S.* Typhi residing in the gallbladder are under sustained immune pressure. The observation that two gallbladder isolates lacked genes encoding proteins for Vi capsule biosynthesis again suggests that these were subject to selective pressure and that the loss of Vi may be an adaptive mechanism for long-term survival. The generation of Vi-negative *S*. Typhi may also question the possibility of their proliferation following mass immunization with TCV.

Identifying genes under selection among gallbladder isolates is crucial for understanding the evolutionary forces and bacterial adaptation to the gallbladder environment during carriage. Signatures of positive selection were detected in a number of genes containing differing gallbladder-specific NS mutations in at least two phylogenetically unlinked gallbladder isolates. Many of these genes are associated with gene regulation under stress and virulence gene expression. For example, the global regulatory gene *rpoS* is responsible for general stress responses and nutrient starvation, and regulates biofilm formation, colonization of Peyer’s patches, persistence in the spleen and the synthesis of Vi ^39–41^. The *degS* gene is involved in salt stress responses and growth-limiting conditions; STY1242 (*ptsG* - glucose-specific PTS system IIBC component) is activated during carbon starvation. These observations suggest that *S.* Typhi is exposed to a range of differing stressors within the gallbladder. Furthermore, the genes responsible for LPS biosynthesis had additionally accumulated NS mutations. LPS is the major component of the outer membrane of Gram-negative bacteria and represents one of the main factors contributing to bile salt resistance ^42,43^. LPS is also a key structural component of the biofilm extracellular matrix forming on human gallstones ^19^. The disruption of genes involved in LPS biosynthesis of *S.* Typhimurium may have a negative influence on biofilm production and attachment ^44^. The enrichment of NS mutations in genes involved in LPS biosynthesis and modification will lead to structural changes in LPS, which we predict will enhance bile resistance and biofilm formation.

This study has its limitations. The number of gallbladder and acute isolates was relatively small and thus might affect the interpretation of the phylogenetic distances between some of the gallbladder isolates and their nearest neighbour. Specifically, our ability to infer associations with uncommon genotypes was limited. Additionally, the identified phylogenetic signature inferred to be associated with carriage was not observed for all gallbladder isolates, due to an underrepresentation in the acutely infected population. Additionally, it was impossible to determine the duration of carriage to confirm our findings, as most typhoid carriers from our study do not recall a history of typhoid ^13^. However, our data suggest that the potential duration of carriage within our gallbladder isolates was variable, which led to variable terminal branch lengths. Despite these limitations, our study is unique and opens up new possibilities for evaluating associations between gallbladder-specific genetic variation and phenotypic differences to better understand the biology of this infectious disease paradox.

## Conclusions

We conclude that typhoid carriage is not associated with any specific genotype nor driven by AMR phenotypes. However, we show that long-term gallbladder carriage results in atypically long phylogenetic branch lengths that can be used to distinguish between carriage and acute infection. Additionally, we found evidence that typhoid carriers are unlikely to play a major role in disease transmission in endemic settings such as Kathmandu, and long-cycle transmission is the primary driver of disease transmission in highly endemic settings. Public health efforts should continue to focus on providing people with safe water and promoting safe food handling and the introduction of TCV to interrupt environmental transmission in endemic settings. It remains important to further investigate the epidemiology, genomics, biology and public health impacts of carriage in parallel to the deployment of these public health measures. The role of carriers may become increasingly important as we move toward eradication, especially as immune selection appears to play a critical role in gallbladder colonisation.

## Methods

### Sampling

Between June 2007 and October 2010, we conducted a *Salmonella* carriage study at Patan Hospital in Kathmandu^13^. In brief, patients undergoing cholecystectomy for acute or chronic cholecystitis were enrolled; bile and stool samples from these patients were subjected to microbiological examination. *S*. Typhi were isolated from bile samples taken from these patients (referred to as gallbladder isolates). Additionally, *S*. Typhi isolates recovered from patients with acute typhoid fever living in the same population recruited into a randomized controlled trial were used for a comparison ^24^ (referred to as acute isolates) (Table S1).

### Bacterial isolation and antimicrobial susceptibility testing

Bile and stool were collected from all cholecystectomy patients for culture. Bile was inoculated into equal volumes of Selenite F broth and Peptone broth and incubated at 37°C overnight. Broth was subcultured onto MacConkey agar and Xylene Lysine Deoxycholate (XLD) agar. After overnight incubation at 37°C, the plates were examined for the growth of Gram-negative bacteria and colonies were identified by API20E (bioMerieux, France). *S*. Typhi were confirmed by slide agglutination using specific antisera (Murex Biotech, Biotech, England).

For the acute isolates, 5-10 ml of blood was taken from all patients with a clinical suspicion of typhoid fever and inoculated into media containing tryptone soya broth and sodium polyanethol sulphonate (up to 25mL). Blood culture bottles were incubated for up to seven days, with blind sub-cultures at 24 hours, 48 hours, and 7 days, or when the broth became cloudy on sheep blood, chocolate, and MacConkey agar. Presumptive *Salmonella* colonies were identified as above.

Antimicrobial susceptibility testing was performed by the modified Bauer-Kirby disc diffusion method with zone size interpretation based on CLSI guidelines ^45^. Etests^®^ were used to determine MICs following the manufacturer’s recommendations (bioMérieux, France). Ciprofloxacin MICs were used to categorise *S*. Typhi isolates as susceptible (≤0.06 μg/mL), intermediate (0.12-0.5 μg/mL) and resistant (≥1 μg/mL) following CLSI guidelines ^45^.

### Vi agglutination assay

Two gallbladder isolates of *S*. Typhi (GB003 and GB428) that lacked the Vi polysaccharide biosynthesis (*viaB*) operon were grown on LB agar plates supplemented with increasing concentrations (1mM, 85mM and 170mM) of NaCl. Vi agglutinations were performed on microscope slides by mixing 10μl of single colony suspensions with 50μl of Vi antisera (Murex Biotech, Biotech, England). Agglutination was recorded after gently agitating the slide for 1 minute. Two gallbladder isolates of *S*. Typhi (GB125 and GB169) containing the *viaB* operon were used as controls.

### Whole genome sequencing and SNP analyses

Total genomic DNA from acute and gallbladder *S*. Typhi isolates was extracted using the Wizard Genomic DNA Extraction Kit (Promega, Wisconsin, USA) (Table S1). 50ng of genomic DNA was subjected to library preparation using the Nextera DNA library prep kit; whole genome sequencing (WGS) was performed on an Illumina MiSeq platform following the manufacturer’s recommendations to generate 250bp paired end reads.

Single nucleotide polymorphisms (SNPs) were called using previously described methods^46^. Briefly, all reads were mapped to the reference sequence of *S.* Typhi strain CT18 (Accession no: AL513382), plasmid pHCM1 (AL513383) and pHCM2 (AL513384) using SMALT (version 0.7.4). Candidate SNPs were called against the reference sequence using SAMtools and filtered with a minimal mapping quality of 30 and a quality ratio cut-off of 0.75. The allele at each locus in each isolate was determined by reference to the consensus base in that genome. This process was performed using *samtools mpileup* and by removing low confidence alleles with consensus base quality ≤20, read depth ≤5 or heterozygous base calls. SNPs in phage regions, repetitive sequences or recombinant regions were excluded, ^47,48^ which resulted in a final set of 2,186 chromosomal SNPs. SNPs were subsequently annotated using the parseSNPTable.py script in the RedDog pipeline (https://github.com/katholt/RedDog). From the identified SNPs in *S.* Typhi genomes, a subset of 68 were used to assign *S.* Typhi isolates to previously defined lineages according to the existing extended *S*. Typhi genotyping framework ^49^.

To identify the potential function of genes containing key SNPs, we investigated the known or predicted functions of the identified genes. We identified SNPs occurring exclusively in either acute or gallbladder isolates and genes containing these SNPs were grouped by their predicted or known function based on the *S.* Typhi functional classification scheme developed by the Sanger Institute (www.sanger.ac.uk) using the genome annotation of *S*. Typhi CT18 ^50^.

The antimicrobial resistance (AMR) gene and plasmid contents of *S*. Typhi isolates were determined using a local assembly approach with ARIBA (Antimicrobial Resistance Identifier by Assembly) ^51^. Resfinder ^52^ and Plasmidfinder ^53^ were used as reference databases of antimicrobial resistance genes and plasmid replicons, respectively.

### Phylogenetic analyses and pairwise SNP distance

A maximum likelihood phylogenetic tree was reconstructed from the SNP alignment of 120 *S*. Typhi isolates (an *S*. Paratyphi A isolate was included as an outgroup) using RAxML (version 8.2.8) with the generalized time-reversible model and a Gamma distribution to model the site-specific rate variation (GTR+Г). Support for the maximum likelihood (ML) tree was assessed via bootstrap analysis with 1,000 pseudoreplicates. Pairwise phylogenetic distances depicting the phylogenetic branch length separating each pair of taxa within subclade 4.3.1 (H58) were estimated using the function *cophenetic* in the ape package (v4.1) in R (v3.3.2). Phylogenetic distances between each taxon and its nearest neighbour on the phylogenetic tree of subclade 4.3.1 were plotted using ggplot2. To investigate the phylogenetic structure of acute and gallbladder *S*. Typhi isolates from Nepal in the global context, a second maximum likelihood tree was inferred from a separate alignment of 23438 SNPs identified from 120 Nepali *S*. Typhi along with 1820 globally representative *S*. Typhi described previously ^54^. A *S*. Paratyphi A isolate was included as an outgroup to root the tree. Support for this ML tree was assessed via 100 bootstrap replicates.

Pairwise genetic distances (the difference in the number of SNPs) within and between acute and gallbladder *S*. Typhi isolates were estimated from the SNP alignment using the ape (v4.1) and adegenet (v2.0.1) packages in R (v3.3.2). Pairwise SNP distances were extracted and plotted using the function *pairDistPlot* in the adegenet package. The Wilcoxon rank-sum test was used for testing the difference in the average pairwise SNP distances between groups.

## Declarations

### Ethics approval and consent to participate

This study was conducted according to the principles expressed in the Declaration of Helsinki and was approved by the institutional ethical review boards of Patan Hospital, The Nepal Health Research Council and The Oxford University Tropical Research Ethics Committee (OXTREC, Reference number: 2108). All enrollees were required to provide written informed consent for the collection and storage of all samples and subsequent data analysis. In the case of those under 18 years of age, a parent or guardian was asked to provide written informed consent.

### Consent for publication

Consent for publication was incorporated as a component of entrance into the study.

### Availability of data and materials

The raw sequence data generated from this study are available in the European Nucleotide Archive (ENA) under the accession numbers described in Table S1.

### Authors’ contributions

Conceptualization: SB

Formal analysis: PTD, NTVT, NTNT, MAR Provided samples: SD, AK, BB

Methodology: PTD, NTVT, NTNT, HNDT, SD, AK, MC, MAR

Writing original draft: DPT, MAR, SB Review and editing: DPT, MC, GD, MAR, SB

Read and approved final version of manuscript: PDT, NTVT, NTNT, HNDT, SD, AK, MC, BB, GD, MAR, SB

## Acknowledgments

We wish to acknowledge all members of the enteric infections group at Oxford University Clinical Research Unit (OUCRU) in Vietnam and Nepal and the study team at Patan Hospital.

## References

1 Crump JA, Crump JA, Luby SP, Luby SP, Mintz ED, Mintz ED. The global burden of typhoid fever. Bull World Health Organ 2004; 82: 346–53.

2 Schwartz E. Typhoid and Paratyphoid Fever. Trop Dis Travel 2010; 366: 144–53.

3 Ledingham JCG. Mr N the milker, and Dr Koch’s concept of the healthy carrier. Lancet 1999; 353: 1354–6.

4 Carrier of typhoid fevver-1912. ; i.

5 Parry CM, Hien TT, Dougan G, et al. Typhoid fever. N Engl J Med 2002; 347: 1770–82.

6 Gonzalez-Escobedo G, Marshall JM, Gunn JS. Chronic and acute infection of the gall bladder by Salmonella Typhi: understanding the carrier state. DOI: 10.1038/nrmicro2490.

7 Prouty AM, Schwesinger WH, Gunn JS. Biofilm formation and interaction with the surfaces of gallstones by Salmonella spp. Infect Immun 2002; 70: 2640–9.

8 Basnyat B, Baker S. Typhoid carriage in the gallbladder. Lancet. 2015; 386: 1074.

9 Schiøler H, Christiansen ED, Høybye G, Rasmussen SN, Greibe J. Biliary calculi in chronic *Salmonella* carriers and healthy controls: a controlled study. Scand J Infect Dis 1983; 15: 17–9.

10 Marshall JM, Flechtner AD, La Perle KM, Gunn JS. Visualization of extracellular matrix components within sectioned Salmonella biofilms on the surface of human gallstones. PLoS One 2014; 9. DOI: 10.1371/journal.pone.0089243.

11 Crawford RW, Rosales-Reyes R, Ramirez-Aguilar M d. l. L, Chapa-Azuela O, Alpuche-Aranda C, Gunn JS. Gallstones play a significant role in Salmonella spp. gallbladder colonization and carriage. Proc Natl Acad Sci 2010; 107: 4353–8.

12 Gonzalez-Escobedo G, Gunn JS. Gallbladder epithelium as a niche for chronic salmonella carriage. Infect Immun 2013; 81: 2920–30.

13 Dongol S, Thompson CN, Clare S, et al. The Microbiological and Clinical Characteristics of Invasive Salmonella in Gallbladders from Cholecystectomy Patients in Kathmandu, Nepal. PLoS One 2012; 7. DOI: 10.1371/journal.pone.0047342.

14 Levine MM, Black RE, Lanata C. Precise estimation of the numbers of chronic carriers of Salmonella typhi in Santiago, Chile, an endemic area. J Infect Dis 1982; 146: 724–6.

15 Mian MF, Pek EA, Chenoweth MJ, Coombes BK, Ashkar AA. Humanized mice for Salmonella typhi infection: new tools for an old problem. Virulence 2011; 2: 248–52.

16 Mateen MA, Saleem S, Rao PC, Reddy PS, Reddy DN. Ultrasound in the diagnosis of Typhoid fever. Indian J Pediatr 2006; 73: 681–5.

17 Mathur R, Oh H, Zhang D, et al. A mouse model of salmonella typhi infection. Cell 2012; 151: 590–602.

18 Crawford RW, Gibson DL, Kay WW, Gunn JS. Identification of a bile-induced exopolysaccharide required for salmonella biofilm formation on gallstone surfaces. Infect Immun 2008; 76: 5341–9.

19 Marshall JM, Flechtner AD, La Perle KM, Gunn JS, Heuser J. Visualization of Extracellular Matrix Components within Sectioned Salmonella Biofilms on the Surface of Human Gallstones. PLoS One 2014; 9: e89243.

20 Yap KP, Gan HM, Teh CSJ, et al. Genome sequence and comparative pathogenomics analysis of a salmonella enterica serovar typhi strain associated with a typhoid carrier in Malaysia. J. Bacteriol. 2012; 194: 5970–1.

21 Baddam R, Kumar N, Shaik S, Lankapalli AK, Ahmed N. Genome dynamics and evolution of Salmonella Typhi strains from the typhoid-endemic zones. Sci Rep 2014; 4. DOI: 10.1038/srep07457.

22 Ong SY, Pratap CB, Wan X, et al. The Genomic Blueprint of Salmonella enterica subspecies enterica serovar Typhi P-stx-12. Stand Genomic Sci 2013; 7: 483–96.

23 Baddam R, Kumar N, Shaik S, et al. Genome sequencing and analysis of Salmonella enterica serovar Typhi strain CR0063 representing a carrier individual during an outbreak of typhoid fever in Kelantan, Malaysia. Gut Pathog 2012; 4: 20.

24 Koirala S, Basnyat B, Arjyal A, et al. Gatifloxacin versus ofloxacin for the treatment of uncomplicated enteric fever in Nepal: an open-label, randomized, controlled trial. PLoS Negl Trop Dis 2013; 7: e2523.

25 Virlogeux I, Waxin H, Ecobichon C, Popoff MY. Role of the viaB locus in synthesis, transport and expression of Salmonella typhi Vi antigen. Microbiology 1995; 141: 3039–47.

26 Steinsiek S, Bettenbrock K. Glucose transport in Escherichia coli mutant strains with defects in sugar transport systems. J Bacteriol 2012; 194: 5897–908.

27 Hatta M, Pastoor R, Scheelbeek PFD, et al. Multi-locus variable-number tandem repeat profiling of Salmonella enterica serovar Typhi isolates from blood cultures and gallbladder specimens from Makassar, South-Sulawesi, Indonesia. PLoS One 2011; 6. DOI: 10.1371/journal.pone.0024983.

28 Baker S, Holt K, Van De Vosse E, et al. High-throughput genotyping of Salmonella enterica serovar Typhi allowing geographical assignment of haplotypes and pathotypes within an urban district of Jakarta, Indonesia. J Clin Microbiol 2008; 46: 1741–6.

29 Holt KE, Baker S, Dongol S, et al. High-throughput bacterial SNP typing identifies distinct clusters of Salmonella Typhi causing typhoid in Nepalese children. BMC Infect Dis 2010; 10: 144.

30 Baker S, Holt KE, Clements ACA, et al. Combined high-resolution genotyping and geospatial analysis reveals modes of endemic urban typhoid fever transmission. Open Biol 2011; 1: 110008.

31 Holt KE, Dolecek C, Chau TT, et al. Temporal Fluctuation of Multidrug Resistant Salmonella Typhi Haplotypes in the Mekong River Delta Region of Vietnam. PLoS Negl Trop Dis 2011; 5: e929.

32 Holt KE, Thomson NR, Wain J, et al. Pseudogene accumulation in the evolutionary histories of Salmonella enterica serovars Paratyphi A and Typhi. BMC Genomics 2009; 10: 36.

33 McClelland M, Sanderson KE, Clifton SW, et al. Comparison of genome degradation in Paratyphi A and Typhi, human-restricted serovars of Salmonella enterica that cause typhoid. Nat Genet 2004; 36: 1268–74.

34 Felix A. Detection of chronic typhoid carriers by agglutination tests. Lancet 1938; 232: 738–41.

35 Nolan CM, White PC, Feeley JC, Brown SL, Hambie EA, Wong KH. Vi serology in the detection of typhoid carrierS. Lancet 1981; 317: 583–5.

36 House D, Ho VA, Diep TS, et al. Antibodies to the Vi capsule of Salmonella Typhi in the serum of typhoid patients and healthy control subjects from a typhoid endemic region. J Infect Dev Ctries 2008; 2: 308–12.

37 Gupta A, My Thanh NT, Olsen SJ, et al. Evaluation of community-based serologic screening for identification of chronic Salmonella Typhi carriers in Vietnam. Int J Infect Dis 2006; 10: 309–14.

38 Lanata CF, Ristori C, Jimenez L, et al. Vi serology in detection of chronic salmonella typhi carriers in an endemic area. Lancet 1983; 322: 441–3.

39 Coynault C, Robbe-Saule V, Norel F. Virulence and vaccine potential of Salmonella typhimurium mutants deficient in the expression of the RpoS (σ(S)) regulon. Mol Microbiol 1996; 22: 149–60.

40 Nickerson CA, Curtiss R. Role of sigma factor RpoS in initial stages of Salmonella typhimurium infection. Infect Immun 1997; 65: 1814–23.

41 Santander J, Wanda S-Y, Nickerson CA, Curtiss R, III. Role of RpoS in fine-tuning the synthesis of Vi capsular polysaccharide in Salmonella enterica serotype Typhi. Infect Immun 2007; 75: 1382–92.

42 Prouty AM, Van Velkinburgh JC, Gunn JS. Salmonella enterica serovar typhimurium resistance to bile: Identification and characterization of the tolQRA cluster. J Bacteriol 2002; 184: 1270–6.

43 Gunn JS. Mechanisms of bacterial resistance and response to bile. Microbes Infect 2000; 2: 907–13.

44 Kim SH, Wei CI. Molecular characterization of biofilm formation and attachment of Salmonella enterica serovar typhimurium DT104 on food contact surfaces. J Food Prot 2009; 72: 1841–7.

45 CLSI. Performance Standards for Antimicrobial Susceptibility Testing. Informational Supplement. 2014 DOI: 10.1186/1476-0711-9-23.

46 Thanh DP, Karkey A, Dongol S, et al. A novel ciprofloxacin-resistant subclade of h58. Salmonella typhi is associated with fluoroquinolone treatment failure. Elife 2016; 5. DOI: 10.7554/eLife.14003.

47 Roumagnac P, Weill F-X, Dolecek C, et al. Evolutionary history of Salmonella typhi. Science 2006; 314: 1301–4.

48 Holt KE, Parkhill J, Mazzoni CJ, et al. High-throughput sequencing provides insights into genome variation and evolution in Salmonella Typhi. Nat Genet 2008; 40: 987–93.

49 Wong VK, Baker S, Connor TR, et al. An extended genotyping framework for Salmonella enterica serovar Typhi, the cause of human typhoid. Nat Commun 2016; 7: 12827.

50 Parkhill J, Dougan G, James KD, et al. Complete genome sequence of a multiple drug resistant Salmonella enterica serovar Typhi CT18. Nature 2001; 413: 848–52.

51 Hunt M, Mather AE, Sánchez-Busó L, et al. ARIBA: rapid antimicrobial resistance genotyping directly from sequencing reads. bioRxiv 2017; : 1–21.

52 Zankari E, Hasman H, Cosentino S, et al. Identification of acquired antimicrobial resistance genes. J Antimicrob Chemother 2012; 67: 2640–4.

53 Carattoli A, Zankari E, Garciá-Fernández A, et al. In Silico detection and typing of plasmids using plasmidfinder and plasmid multilocus sequence typing. Antimicrob Agents Chemother 2014; 58: 3895–903.

54 Wong VK, Baker S, Pickard DJ, et al. Phylogeographical analysis of the dominant multidrug-resistant H58 clade of Salmonella Typhi identifies inter- and intracontinental transmission events. Nat Genet 2015; 47: 632–9.

